# A Bioinformatic Analysis of BAG Protein Interactors and Pathways in Alzheimer’s and Parkinson’s Disease

**DOI:** 10.1101/2025.07.18.665492

**Authors:** Sudarshan Ramanan, Gail VW Johnson

## Abstract

Alzheimer’s disease (AD) and Parkinson’s disease (PD) are the two most common neurodegenerative disorders. While the symptoms and general etiology may be different, these two diseases share significant common features in terms of their disease pathogenesis. Within the scope of neurodegenerative disorders, the Bcl-2 associated athanogene (BAG) family proteins and associated interactors have been a key area of focus. The BAG family is a group of proteins that contain at least one evolutionarily conserved BAG domain. Despite this similarity, their interactions and functions can vary widely. So far, research has predominantly scrutinized individual BAG proteins, rather than explore potential cooperative actions among family members. Some BAG family members may function together thereby indicating potential interactions within this family. Although connections among BAG members have been observed, their role in neurodegenerative disorders, such as AD and PD, remains largely uncharacterized. This mini review explores the common pathways, intersections, and differences within these interactions as well as their link to AD and PD. Using computational techniques to mine transcriptomic data, several groupings of pathways that these BAG family members are involved in were identified in the context of AD and PD. Understanding these pathways and their relationships may uncover potential gaps in current research and help identify novel therapeutic targets for the treatment of these neurodegenerative diseases.

**Significance statement:** Although distinct diseases, Alzheimer’s disease and Parkinson’s disease share common features such as protein aggregation and mitochondrial dysfunction. Members of the BAG family of proteins have been implicated in the pathogenesis of both diseases. Computational techniques were used to mine transcriptomic data of Alzheimer’s and Parkinson’s disease cases to identify common pathways. BAG protein interactors, common to all family members, were analyzed in the context of these common pathways for Alzheimer’s and Parkinson’s disease. These analyses provide insights into the pathways mediated by these BAG protein interactors that are likely at the intersection of Alzheimer’s disease and Parkinson’s disease pathologies.

## Introduction

Alzheimer’s disease (AD) is characterized by amyloid plaques and neurofibrillary tangles (NFTs) that result from the accumulation of predominantly extracellular Aβ and intracellular tau, respectively (Nelson et al., 2009). In the disease state, the processing of amyloid precursor protein (APP) results in increases in Aβ40 and Aβ42, which favors the formation of neurotoxic oligomers (Gu and Guo, 2013). These oligomers often form clusters around cerebral vessels and accumulate predominantly in the neocortex in AD brains (Zhang et al., 2023). NFTs, on the other hand, are formed by the aggregation of tau polymers (Binder et al., 2005). In non-AD brains, tau is bound to microtubules and is phosphorylated by kinases, which allows for neuronal plasticity and structural changes (Mietelska-Porowska et al., 2014). A dysregulation of this process, however, can lead to the hyperphosphorylation of tau, which contributes to the aggregation of tau into oligomers and NFTs (Ward et al., 2012). The oligomers are likely the more toxic pathological species as they have been reported to disrupt synaptic function and axonal transport mechanisms (Singh et al., 2020; Swanson et al., 2017). As a result of these dysregulated processes, AD patients experience cognitive decline and progressive dementia.

The second most common neurodegenerative disease, Parkinson’s Disease (PD), is characterized by the gradual degeneration of midbrain dopamine neurons in the substantia nigra (Aarsland et al., 2021). Reactive oxygen species produced by enzymatic and non-enzymatic reactions result in the oxidation of dopamine, which contributes to neurodegeneration in PD (Naoi and Maruyama, 1999; Segura-Aguilar et al., 2014). The loss of dopaminergic neurons is a key contributor to the tremors, rigidity, and bradykinesia that are hallmark symptoms of PD. Furthermore, the presence of Lewy bodies, which are composed primarily of α-synuclein, is a key neuropathological aspect of PD (Mensikova et al., 2022). Abnormal forms of α-synuclein (protofibrils) are a significant driver of progressive neuronal death; impairing mitochondria, disrupting lysosomal function, and altering calcium homeostasis (Stefanis, 2012). Despite this knowledge, the underlying molecular mechanisms of PD remain incompletely understood.

When studying AD and PD, it is important to examine some of the known crossovers between these two diseases. For example, oxidative damage to coding and non-coding RNA is known to result in gene dysregulation, a pathological outcome often observed in both AD and PD (Nunomura and Perry, 2020). Further, the most prominent genetic risk factor for AD is the ε4 allele of the apolipoprotein E (APOE) gene, located on chromosome 19, which has also been linked to many PD cases (Giau et al., 2015; Szwedo et al., 2022). While the pathophysiologic similarities between AD and PD may serve as a microcosm of the wider neurodegenerative disorder spectrum, focusing on the shared interactors of these two disorders may prove to be an important therapeutic target. This is where the B-cell lymphoma 2 (Bcl-2)-associated athanogene (BAG) family of proteins may provide insights into the underlying pathogenic processes at play.

### B-cell lymphoma 2 (Bcl-2)-associated athanogene (BAG) Family

The BAG family is a group of multi-functional proteins that play a crucial role in various biological processes including maintaining cellular homeostasis, regulating cell proliferation, and controlling apoptosis (Qian et al., 2022). Characterized by the presence of at least one BAG domain located in the C-terminal region of their structure, BAG proteins belong to an evolutionary conserved family (Jiang et al., 2023; Kabbage and Dickman, 2008). The BAG domain of these proteins interacts with the nucleotide binding domains (NBDs) of Heat Shock Protein 70 kDa (Hsp70), encoded by the HSPA1A gene, and Heat Shock Cognate (Hsc70), which is encoded by the HSPA8 gene, to regulate their chaperone activity (Rauch et al., 2016; Perez-Vargas et al., 2006). This enables the BAG proteins to function as nucleotide exchange factors (NEFs), facilitating the release of adenosine diphosphate (ADP), which in turn permits adenosine triphosphate (ATP) to bind with HSPA1A/HSPA8 (Bracher and Verghese, 2015). This interaction also modulates other processes involved in mediating apoptosis, signal transduction, and cell proliferation (Rasola et al., 2001). Despite the fact that they all contain a well-defined BAG domain, each individual BAG protein also performs distinct functions (Qin et al., 2016). It is important to note that based on the available literature, all these BAG proteins, with the exception of BAG4, have been reported to have specific functions in the central nervous system (CNS). Given the lack of reports on BAG4’s involvement in the CNS, it will not be included in future discussions.

### BAG1

There are four major isoforms of the BAG1 gene due to alternative translation initiation from a single mRNA transcript (Knee et al., 2001). As a group, these isoforms are especially important for their function in protein quality control mechanisms through the formation of a carboxyl terminus of the HSPA8-interacting protein (CHIP)-HSPA1A-BAG1 complex (Hantouche, 2017; Quintana-Gallardo, 2019). This is of particular importance since CHIP is an E3 ligase that acts as a co-chaperone to aid in the refolding of misfolded proteins and promotes the degradation of misfolded proteins via proteasomal pathways (Shankar et al., 2024). Furthermore, in addition to the NBD of HSPA1A, BAG1’s BAG domain interacts with the mitogen-activated protein kinase (MAPK) Raf-1, inducing its activation (Suyama et al., 2023).

BAG1 plays a key role in inhibiting apoptosis (Cutress et al., 2002). BAG1 achieves this in part by binding to and forming a complex with anti-apoptotic proteins such as Bcl-2 and MCL1 that prevent pro-apoptotic proteins from initiating cellular death (Aveic et al., 2011). As a result of the interaction of BAG1 with Bcl-2 or MCL1, these proteins translocate to the mitochondria, which is crucial in preventing mitochondrial outer membrane permeabilization (MOMP) (Kalkavan and Green, 2018; Popgeorgiev et al., 2018). Downstream, MOMP allows the release of cytochrome c and other pro-apoptotic proteins to promote caspase activation and apoptotic cell death (Garrido et al., 2006). Given its role in inhibiting apoptosis, it is not surprising that knockout of BAG1 in mice results in embryonic lethality due to widespread apoptosis in the developing nervous system (Inose-Maruyama et al., 2024; Götz et al., 2005). In terms of BAG1’s neuroprotective effects, overexpression of BAG1 reduces injury after a stroke and promotes axonal outgrowth after an optic nerve crush (Planchamp et al., 2008; Kermer, 2003). This indicates that BAG1 is an important player in maintaining cellular and protein homeostasis.

### BAG2

BAG2, although it is a bona fide member of the BAG family, has a C-terminal BAG domain that exhibits low homology with other BAG domains. In fact, this domain has often been referred to as the “brand new BAG” (BNB) domain. Nonetheless, this “BNB” domain functions as a BAG domain as it binds HSPA1A/HSPA8 and regulates nucleotide exchange and ATP binding (Chen et al., 1996; Xu et al., 2008). Studies have even shown BAG2 to have a relatively high affinity for the NBD of HSPA1A (Takayama et al., 1999).

BAG2, like BAG1, plays a key role in the degradation of misfolded proteins, through its interaction with CHIP, an E3 ligase involved in regulating protein quality control. However, in contrast to BAG1, BAG2 inhibits the E3 ubiquitin ligase activity of CHIP by decreasing CHIP’s association with the ubiquitin-conjugating enzymes E2 and thus inhibits ubiquitylation of clients allowing for chaperone-assisted maturation (Arndt et al., 2005). BAG2 also efficiently inhibits the CHIP-mediated ubiquitylation of HSPA1A by preventing its interaction with E2 (Schonbuhler et al., 2016). Interestingly, in response to specific forms of stress, where refolding is not possible, BAG2 facilitates the formation of protein condensates. BAG2 then directs these phase-separated membrane-less organelles to 20S proteasomes resulting in ubiquitin-independent client degradation. (Carrettiero et al., 2022). This allows BAG2 to help a cell maintain proteostasis by facilitating ubiquitin independent proteasome degradation of client proteins when refolding is not possible (Carrettiero et al., 2022).

### BAG3

In addition to its C-terminus BAG domain, BAG3 contains a WW domain near its amino terminus and a proline-rich repeat (PXXP) region, which helps mediate its binding to other proteins (Iwasaki et al., 2010). Further, the presence of the highly conserved Ile-Pro-Val (IPV) domains in its N-terminus facilitates its interactions with small Hsps such as HspB8 and HspB6 (Cristofani et al., 2019; Fuchs et al., 2009). This can play an important role in the inhibition of protein aggregation (Knezevic et al., 2015). Similar to the other BAG family members, the BAG domain of BAG3 interacts with the NBD of HSPA1A to mediate important processes (Doong et al., 2002). One such process of particular focus in the context of neurodegenerative disorders relates to chaperone-assisted selective autophagy, which is a major intracellular degradation system that converges with the endosomal-lysosomal pathway (Yim and Mizushima, 2020).

BAG3 plays a significant role in regulating the activity of Rab35, a notable small GTPase that facilitates the movement of cellular membranes between the plasma membrane and endosomes in eukaryotic cells (Lin et al., 2022). BAG3 modulates Rab35 activity by regulating TBC1D10B, a member of the RabGAP family, which forms a complex with the activated state of Rab35, facilitating the hydrolysis of GTP (Lin et al., 2022). BAG3 associates with TBC1D10B, which limits its GAP activity towards Rab35 thus allowing Rab35 to remain active. When BAG3 is depleted, this results in a decrease in Rab35 activity, which is restored when TBC1D10B is also depleted, confirming that BAG3 regulates Rab35 through tis interaction with TBC1D10B (Lin et al., 2022). Previous studies have shown that since BAG3 facilitates the clearance of tau, and Rab35 mediates the clearance of specific phosphorylated tau (p-tau) species, it is likely that the BAG3-TBC1D10B-Rab35 signaling axis plays a key role in the clearance of p-tau (Lei et al., 2015; Vaz-Silva et al., 2018). In support of this hypothesis, depletion of TBC1D10B results in increased levels of tau that is phosphorylated at specific AD-relevant epitopes (Lin et al., 2022). Overall, these data strongly suggest that BAG3 acts in concert with the TBC1D10B-Rab35 axis to regulate the clearance of p-tau (Lin et al., 2022). Interestingly, recent data has pointed towards notable reductions in BAG3 levels in AD brains compared to controls (Zhou et al., 2020). Additionally, increased BAG3 expression appears to have protective effects against the progression of tau pathology (tauopathy) and α-synuclein pathology (Sheehan et al., 2023; Fu et al., 2019).

### BAG5

Compared to the other BAGs, BAG5 is considered unique since it comprises five BAG domains in tandem (Gupta et al., 2022; Briknarová et al., 2002). The fifth binding domain is what allows the protein to interact with HSPA1A inducing certain conformational changes in the NBD of HSPA1A, which results in its loss of affinity for ADP binding (Arakawa et al., 2010). By facilitating the release of ADP from HSPA1A and allowing for a new ATP binding cycle, BAG5 exhibits NEF activity by enhancing HSPA1A’s ability to refold misfolded proteins (Fernandez-Fernandez and Valpuesta, 2018).

Within the context of diseases, mutations in the PINK1 and Parkin gene are commonly associated with the pathogenesis of PD (Brooks et al., 2009). Studies have shown that BAG5 can inhibit the activity of Parkin E3 ligase activity and HSPA1A refolding activity leading to an increase in dopaminergic neuron degeneration (Kalia et al., 2004). BAG5 has also been shown to interact with DJ-1 inhibiting its protective effects against mitochondrial oxidative damage (Qin et al., 2017).

### BAG6

BAG6 is the largest member of the BAG family, consisting of approximately 1132 amino acids (Mock et al., 2015). In addition to its amino-terminal UbL domain and carboxy-terminal BAG domain, BAG6 also features a nuclear localization signal (NLS) at its carboxy terminus, a caspase 3 cleavage site, an extended proline-rich region, and a zinc-finger-like domain situated in the central part of its primary protein structure (Lee and Ye, 2013). BAG6 is a key regulator of autophagy as it controls the intracellular localization of different acetyltransferases (Pattingre and Turtoi, 2022; Xu and Wan, 2022). This is especially important in situations where homeostasis is compromised as BAG6 can promote the upregulation of pro-autophagic genes (e.g., SESN1/Sestrin1) through acetylation of the transcription factor p53 and enhanced autophagosome formation (Sebti et al., 2014). BAG6 has been shown to localize in mitochondria, and increased expression of BAG6 induces mitochondrial fission, activates the Pink1/Parkin pathway and induces mitophagy. Interestingly. BAG6 has 2 LC3 interacting regions (LIRs) and the C-terminal LIR is essential for mitophagy induction, suggesting that BAG6 may function as a mitophagy/autophagy receptor (Ragimbeau et al., 2021; Wirth et al., 2019).

### Study Objective

Despite the differences in structure and interactions of the various BAG proteins, most studies focus solely on the role of one or two BAG members and their classical interactors (Bcl-2 or HSPA1A/HSPA8). Studies have suggested, however, that certain BAG family members may work together, indicating potential cooperative interactions within the BAG family (Lin et al., 2025). Still, the functional and cellular significance of these interactions remain poorly understood. Addressing these knowledge gaps is essential to exploring the potential of BAG proteins as a diagnostic tool and therapeutic target in neurodegenerative disorders.

To begin analyzing the interactors and pathways regulated by the BAG protein family members in the context of AD and PD, we first examined the five members of the BAG family and assessed the common biological processes in which they are involved based on their shared interactors. Two different RNA sequencing studies focused on AD and PD, (Cappelletti et al., 2023; Guennewig et al., 2021) were analyzed, and the common biological processes involved in these two disorders were studied. Finally, the functions of the BAG family members were compared with the pathways of neurodegenerative disorders (AD and PD). Identifying such similarities and differences may provide unique insight into the ways the BAG family regulates neurodegenerative disorders.

## Methodology

### Dataset Selection and Data Preprocessing

BioGRID (https://thebiogrid.org) is a publicly accessible repository containing data from protein, chemical, and genetic interactions that have been curated from thousands of publications and databases (Oughtred et al., 2021). Given the role of the aforementioned BAG family members in the progression of neurodegenerative disorders, the corresponding BAG identifiers, BAG1 through BAG6 (excluding BAG4 due to a paucity of data surrounding its role in the CNS), were assessed within the context of the *homo sapiens* species. As part of the study by Lin et al., 2025, a final dataset of approximately 2500 interactors including 476 BAG1 interactors, 623 for BAG2, 634 for BAG3, 343 for BAG5, and 453 interactors for BAG6 was curated.

### Gene Interactors

Corroborated by a recent study Lin et al., 2025, 4 specific interactors common to all BAG proteins (with the exception of BAG4) were identified and were used for further analysis. These interactors include HSPA2, HSPA1A, METTL3, and CHIP (the protein encoded by the gene, STUB1).

### Biological Processes

Enrichr is a tool that compares gene lists to numerous other gene databases that include information of molecular functions, biological processes, and pathways (https://maayanlab.cloud/enrichr/). At a high level, this tool works by comparing a list of genes to a library and mapping those genes based on commonly found processes in experiments and other databases. The output of this would indicate a set of enriched processes and pathways various inputted genes are involved in. Using this approach, the common biological processes of the 4 previously mentioned shared interactors were first identified and then filtered based on adjusted p-value (cutoff of 0.05), which indicates existence of significant pathways or processes. Based on the collective dataset of all BAG proteins, each BAG protein, and its corresponding Entrez gene symbol was analyzed using the Enrichr toolkit with a specific focus on the Ontologies feature. Using the Gene Ontology (GO) Biological Process knowledgebase (2023), several hundred molecular function clusters were identified and rank ordered based on most to least significant (adjusted p-value ≤ 0.05).

### Transcriptomic Analysis

To further assess the roles these aforementioned BAG protein family members, play in neurodegenerative disorders, transcriptomic data from two separate studies were assessed. The focus of this analysis was on AD and PD given their prevalence and similarities in pathological indicators (Caligiore et al., 2022; Nuytemans et al., 2016; Xiang et al., 2024). RNA sequence data from cerebral cortex tissue samples in AD patients (n=22) and control were compared with RNA sequence data from cerebral cortex (Braak Lewy stage 5 versus control) tissue samples in PD patients (n=42) (Cappelletti et al., 2023; Guennewig et al., 2021). Significance was set at a false discovery rate-adjusted p value ≤ 0.05 for both studies.

Upon independently assessing the set of differentially expressed genes in both studies, the corresponding biological processes implicated based on these genes were then examined using GO enrichment terms. A total of 2385 GO terms were associated with AD, while 2796 processes were associated with the PD differentially expressed genes (p-value ≤ 0.05). The overlap of these sets of GO biological processes yielded a final set of 1569 shared biological processes, which served as the basis for the common processes involved in the progression of AD and PD (**Figure 1**). The significantly enriched pathways based on the shared interactors of the BAG protein family members were then compared to the common processes involved in neurodegenerative disorders. This resulted in a final number of 77 overlapping pathways and processes (p-value ≤ 0.05) shared between BAG interactors and AD and PD (**1**).

**Figure 1:**
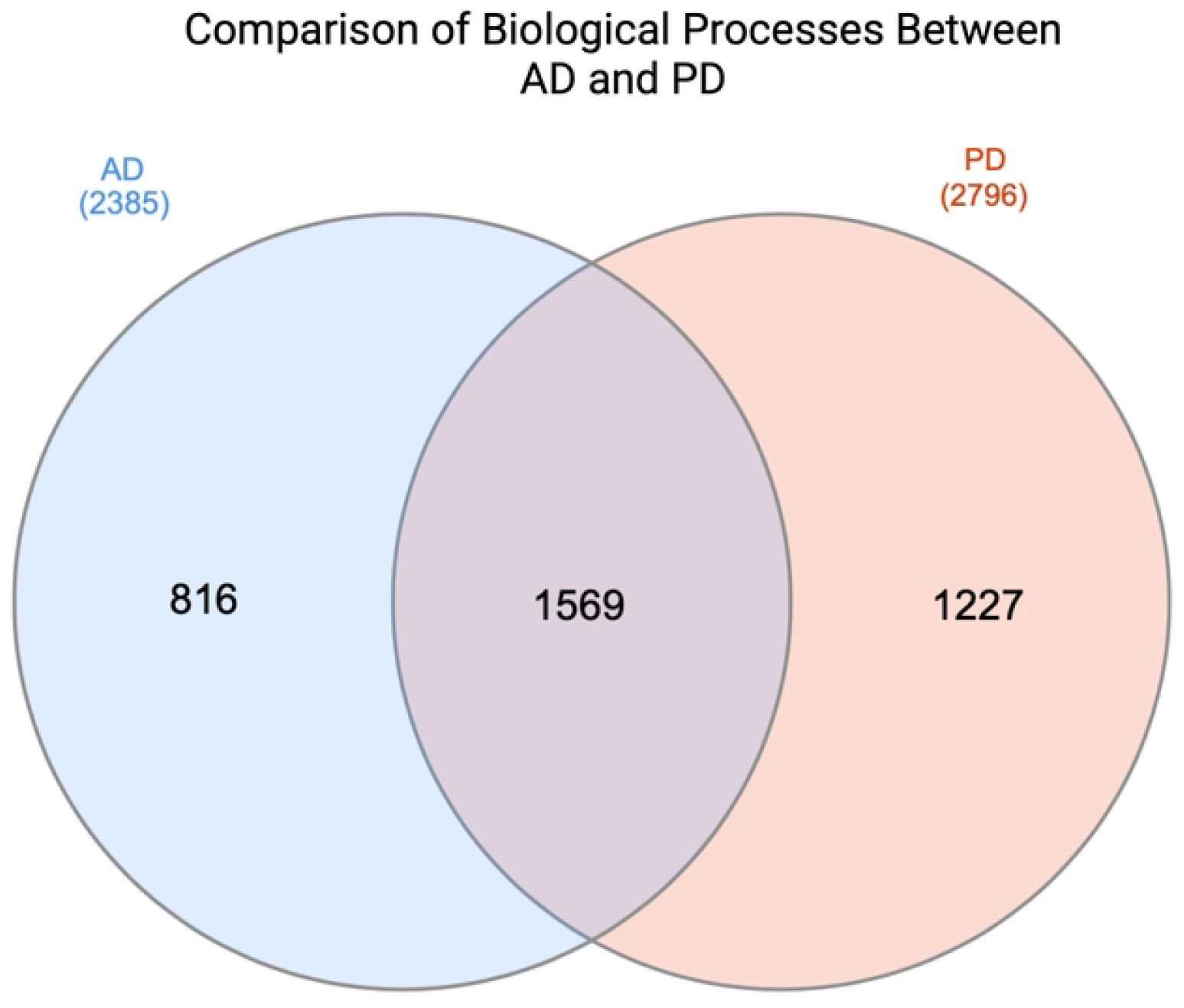
Comparison of common GO biological processes between AD and PD datasets based on the differentially expressed genes (p-value ≤ 0.05). The PD set contains 2796 biological processes based on the differentially expressed genes in disease versus control. The AD set contains 2385 biological processes based on the differentially expressed genes in disease versus control. The intersection (1569) refers to the common processes or overlaps between both sets, while 816 and 1227 refer to the specific processes only involved in either AD or PD respectively. Created with InteractiVenn.

## Results and Discussion

Gaining insight into the key interactions that are common across the different BAG proteins is vital to understanding the potential functional pathways and biological mechanisms implicated in neurodegenerative disorders like AD and PD. A combination of computational and bench-side techniques can help researchers develop a more comprehensive understanding of these diseases. Within the context of AD and PD, and BAG protein family members, computational techniques indicate that 4 primary genes are common interactors across all BAG proteins (Lin et al., 2025). These may also serve as novel targets for therapeutic interventions and warrant further exploration.

The 4 primary interactors identified by Lin et al., 2025 were found upon mining the BioGRID database. The common interactors across all five BAG protein family members were, HSPA2, HSPA1A, CHIP, and METTL3. These 4 interactors may serve as a conduit or bridge among the different BAG proteins potentially indicating an avenue for compensatory mechanisms at play. It is important to note that while these 4 interactors themselves might not be directly involved in AD or PD, they still may play a part in the key biological processes of these neurodegenerative disorders. In addition, these 4 interactors may lie upstream or downstream of the identified differentially expressed genes that are known to be involved in AD and PD (**Figure 1**). Comparing the processes these interactors are involved in, alongside the general processes of AD and PD, could provide important insights into how these BAG domains may specifically regulate neurodegenerative disorders and serve as novel therapeutic targets.

### Importance of Identifying these Interactors

Upon establishing these common (conserved) interactors, it is important to explore the general scope of the biological processes that lie at the intersection of these BAG domains. These 4 BAG interactors are involved in approximately 145 key biological processes. To understand their specificity to certain neurodegenerative disorders, however, it is vital to compare these pathways with those found in AD and PD. Comparing the common processes of the BAG interactors and the processes implicated in neurodegenerative disorders, it becomes clear that there are 77 overlapping biological processes (**Table 1**). These pathways can then be separated into 8 distinct subcategories that have differing roles within the overall regulation of neurodegenerative disorders by BAG proteins (**Table 1**).

**Table 1:**
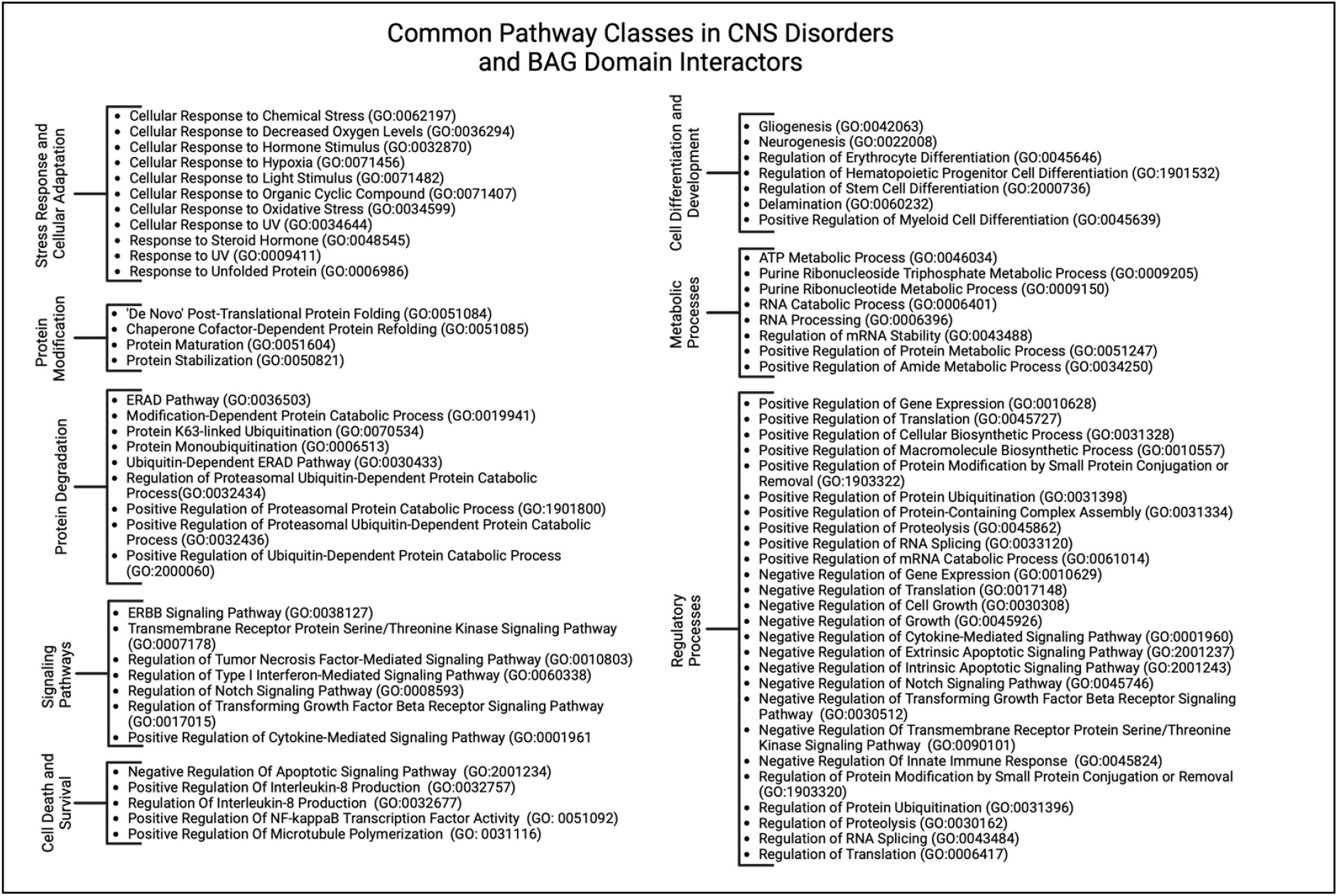
Summary of the main identified classes of common GO biological processes between neurodegenerative disorders and the most significant interactors of the various BAG domains.

### A Deeper Dive into Key Pathways

Out of all the pathway groups identified, protein modification and degradation pathways are especially crucial since they are both characterized by protein misfolding, abnormal posttranslational processing, and protein aggregation in AD and PD (**Table 1**). In neurodegenerative disorders, maintaining homeostasis is of utmost importance when protein-protein interactions are disturbed and the potential for misfolded proteins increases (Ashraf et al., 2014). More specifically, important mechanisms such as the ubiquitin-proteasome system and the autophagy-lysosomal pathway can become severely compromised (Pan et al., 2008). The dysregulation of these pathways can lead to accumulation of toxic tau aggregates. Different members of the Hsp family, including HSPA1A and HSPA2, help ensure that larger protein complexes are not severely altered in diseased states. These various Hsps work together to preserve protein stability and conformation in order to maintain cellular homeostasis, which can be found in the “Protein Stabilization” pathway in **Table 1** (Kumar et al., 2022). Studies have likewise shown that Hsps are transported to synapses and axons and work to block the aggregation of misfolded proteins alongside promoting their degradation (Zatsepina et al., 2021). Further, HSPA2 has been identified as a possible biomarker for AD as it is strongly upregulated in AD cases (Dong et al., 2022). HSPA1A, HSPA2, and HSPA8 are also known to regulate protein misfolding, including inhibition of Aβ aggregation and NFT formation in AD, thereby falling under the general umbrella pathway term of “Chaperone cofactor-dependent protein refolding” in **Table 1** (Dong et al., 2022). In addition, HSPA1A facilitates the disaggregation and degradation of α-synuclein fibrils (Li et al., 2024). HSPA1A and HSPA8 have been suggested to recognize the N-terminus of α-synuclein near the Tyr39 segment thereby preventing its aggregation (Rutledge et al., 2022). Furthermore, inhibiting the interaction between α-synuclein and HSPA8 specifically has been shown to promote aggregate formation, in line with studies that have shown that HSPA1A knockdown can also promote different forms of α-synuclein oligomerization (Sirtori et al., 2020). While reducing both HSPA1A and HSPA8 tends to increase monomeric α-synuclein levels, only HSPA8 knockdown is found to significantly enhance oligomer formation (Sirtori et al., 2020). This effect is likely due to impaired α-synuclein degradation, as HSPA8 plays a more central role in chaperone-mediated autophagy compared to HSPA1A (Sirtori et al., 2020). Similar to the effect on α-synuclein, HspA1A also prevents tau oligomerization and facilitates tau degradation plays (Young et al., 2016; Patterson et al., 2011). Further, recent data has implicated HSPA2 as a specific key regulator of late-onset AD (Petyuk et al., 2018) Overall, it is clear that the Hsp family plays a key role in regulating crucial proteins involved in the progression of AD and PD.

CHIP interacts with members of the Hsp family including HSPA1A (Xu et al., 2008). Due to its E3 ubiquitin ligase activity, CHIP is able to regulate the stability and abundance of misfolded proteins by targeting them for degradation, which can have a significant impact on overall protein turnover (Tawo et al., 2017). This is closely associated with the “Positive regulation of proteasomal protein catabolic process” and “Positive regulation of ubiquitin dependent protein catabolic process” pathways found in **Table 1**. When examining the attributes of CHIP that enable this, it is important to take a closer look into specific domains. The tetratricopeptide repeat (TPR) domain at the N-terminus is essential for CHIP’s interaction with chaperones and other substrates (Tedesco et al., 2023). Likewise, the C-terminal U-box domain serves as a catalytic center for the protein’s E3 ubiquitin ligase activity (Yang et al., 2021). These two specific domains help assemble an important signaling complex involving IFNγ-receptors, which CHIP subsequently destabilizes by targeting IFNγ-R1 and JAK1 for proteasomal degradation (Apriamashvili et al., 2022). Within the context of AD, CHIP has been found to bind more tightly to p-tau than unmodified tau, inhibiting the aggregation of phosphorylated proteoforms of microtubule-associated protein tau (MAPT) (Nadel et al., 2023). In the context of PD, in vivo studies have shown that CHIP can also mediate the degradation of α-synuclein aggregates (Dimant et al., 2014). Age-dependent decreases in CHIP tend to correlate with increased accumulation of NQO1, an enzyme that protects cells from oxidative stress and damage. Interestingly, in approximately half of the AD cases examined in this study, due to polymorphism NQO1 protein levels were undetectable, suggesting that the age-dependent accumulation of NQO1 is impaired in certain AD cases (Tsvetkov et al., 2011). CHIP is required for the degradation of hypoxia-inducible factor-1A (HIF1A) by chaperone mediated autophagy (Ferreira et al., 2013). It is suggested that CHIP, in association with HSPA8 and other co-chaperones, facilitates substrate degradation through both the proteasomal and lysosomal pathways (“Positive regulation of proteasomal protein catabolic process” in **Table 1**) (Tedesco et al., 2023). Given the functions of CHIP, enhancing CHIP expression is being considered as a therapeutic strategy for both AD and PD (Zhang et al., 2020).

The addition of m6A to mRNA by METTL3 is an important modification that can significantly impact protein synthesis (Xia et al., 2023). METTL3 itself plays a key role in what is known as the “m6A writer complex” (Meng et al., 2023). This allows METTL3 to self-interact, which subsequently drives condensation and phase separation (Han et al., 2022). This phase separation in the nucleus enables the formation of a multilayer condensate thereby aiding in ribosomal assembly (Lafontaine et al., 2021). This then helps create the writer complex by forming a stable heterodimer with METTL14, which is critical for substrate recognition (“Modification-dependent protein catabolic process” in **Table 1)** (Zeng et al., 2023). In AD, not only is METTL3 expression decreased in the hippocampus, but in a C57BL/6 mouse model of METTL3 knockdown, it was found that the absence of this gene further exacerbates cognitive deficits (Zhao et al., 2021). This is most likely because knockdown of METTL3 in the hippocampus can lead to synaptic loss and neuronal death, along with AD-related cellular alterations (Zhao et al., 2021). Moreover, the m6A modification, itself, is decreased in AD brains (Zhao et al., 2021). On the other hand, upregulation of METTL3 enhances autophagic clearance of p-tau through m6A-dependent regulation of CHIP. Furthermore, overexpression of METTL3 has been shown to rescue synaptic damage and cognitive impairment in Aβ-induced AD mice (Zhao et al., 2021). Similarly, in PD, a recent study (n=172) found lower mRNA levels of m6A, METTL3, and METTL14 in PD patients compared to healthy controls with METTL14 primarily driving the abnormal m6A modifications (He et al., 2023). In MPTP-induced Parkinson’s disease mouse models, the upregulation of METTL3 by the transcription factor NRF1 increases m6A modifications on the mRNA of glutaredoxin (GLRX). This modification stabilizes the mRNA through the reader protein IGF2BP2, leading to a reduction in oxidative stress and dopaminergic neuron degeneration, ultimately improving motor coordination in behavioral tests (Gong et al., 2024).

### Hierarchical Clustering

While it is important to understand the specific similarities and differences among different interactors, processes, and pathways in various BAG proteins at a granular level, it is equally crucial to take a step back and understand the general relationships among the different BAG proteins. Based on the interactors across all 5 BAG proteins, BAG2 and BAG5 tend to have the greatest similarity to each other and, therefore, cluster together (**Figure 2**). The clustering of BAG2 and BAG5 could indicate a closer evolutionary relationship between these two proteins compared to other BAG family members purely based on interactors. The hierarchical clustering of biological processes, on the other hand, groups BAG1 and BAG5 together, alongside BAG2 and BAG6 (**Figure 2**). Likewise, such clustering could indicate shared functional roles of these proteins within the overarching regulatory and responses network of BAG proteins in neurodegenerative disorders. While these domains may not share genes themselves, the general grouping of genes among these proteins may be involved in similar processes. The direct interactors of BAG2 and BAG5 may have been genetically similar at one point but may have developed distinct biological functions over time (Lin et al., 2025). These proteins may even be involved in a more complex system of interactions that participate in related cellular pathways but may have less genetic similarity. Overall, in certain cellular contexts, these BAG family members may have compensatory mechanisms that can play a key role in the regulation of neurodegenerative disorders.

**Figure 2:**
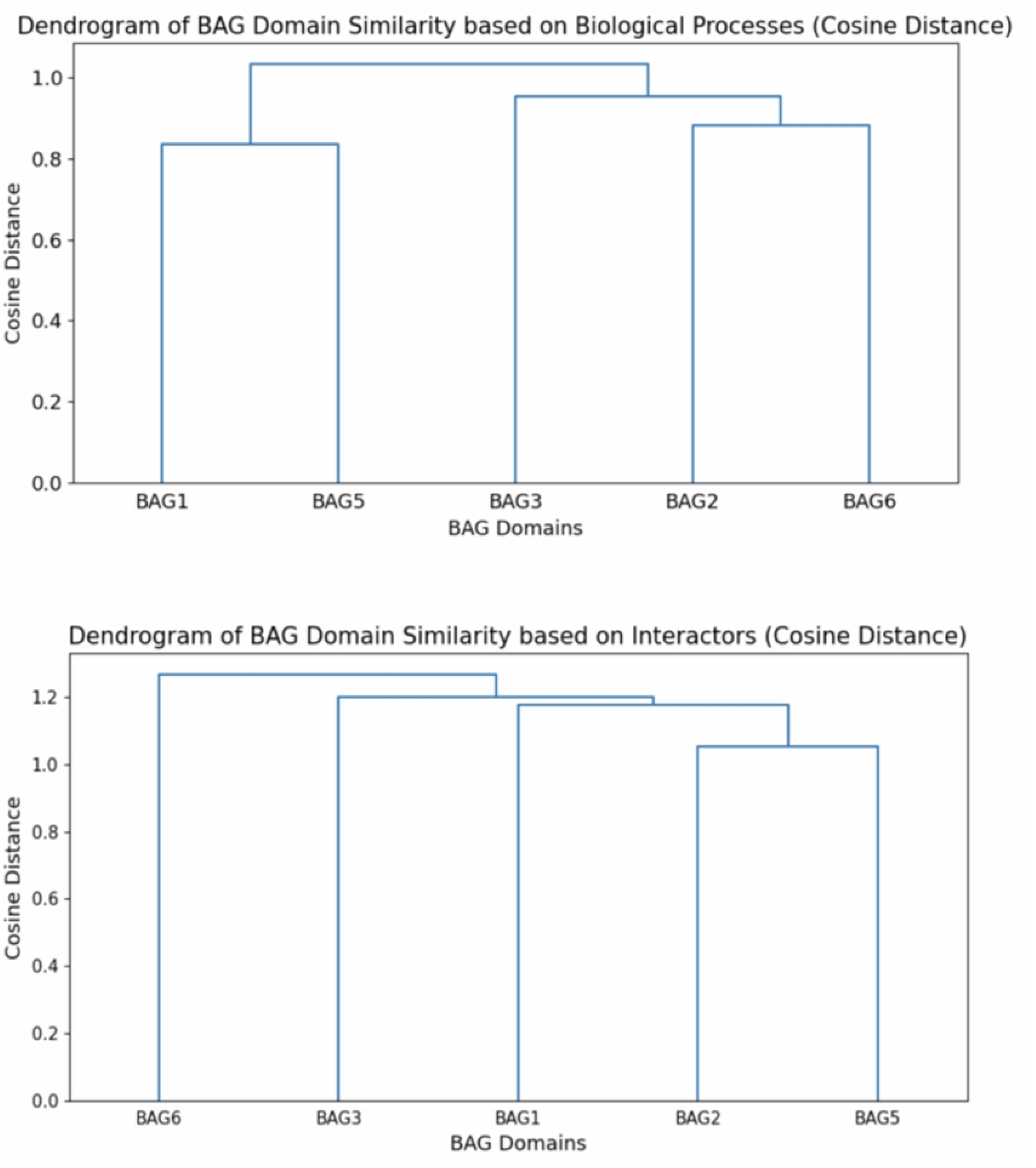
Dendrogram of BAG family members clustered together based on the similarity of biological processes and interactors using the cosine distance metric. Created using Python.

## Conclusion

Though each clustering may be different depending on the specifics of the analysis, these relationships provide insight into which specific BAG proteins may have comparatively stronger or weaker compensatory responses in neurodegenerative disorders. Understanding the relationships of different processes and pathways will provide insights into potential compensatory mechanisms at play (Kim and Suh, 2022). BAG proteins with similar functions and interactors are more likely to compensate for each other’s loss or dysfunction. Understanding the role that the shared BAG interactors play within key pathways, may provide insight into the broader influence of the BAG family in the regulation of AD and PD. The 77 identified pathways not only indicate unique ways the BAG family members interact with the dysregulated systems in AD and PD, but they also provide direction for future research to examine other interactors and mechanisms at play. Furthermore, these pathways also suggest novel therapeutic targets. While this work provides a foundation for understanding the relationships of the BAG protein family, experimental validation would be necessary to confirm these potential compensatory responses and functional similarities.

